# Identifying the effect of vancomycin on HA-MRSA strains using bacteriological and physiological media

**DOI:** 10.1101/2020.05.06.079640

**Authors:** Akanksha Rajput, Saugat Poudel, Hannah Tsunemoto, Michael Meehan, Richard Szubin, Connor A. Olson, Yara Seif, Anne Lamsa, Nicholas Dillon, Alison Vrbanac, Joseph Sugie, Samira Dahesh, Jonathan M. Monk, Pieter C. Dorrestein, Rob Knight, Joe Pogliano, Victor Nizet, Adam M. Feist, Bernhard O. Palsson

## Abstract

Healthcare-associated methicillin-resistant *Staphylococcus aureus* (HA-MRSA) USA100 strains are of major concern due to their evolving antibiotic resistant. They are resistant to a broad class of antibiotics like macrolides, aminoglycosides, fluoroquinolones, and many more. The selection of appropriate antibiotic susceptibility examination media is very important. Thus, we use bacteriological (CA-MHB) as well as physiological (R10LB) media to determine the effect of vancomycin on USA100 strains. The study includes the profiling behaviour of HA-MRSA USA100 D592 and D712 strains in the presence of vancomycin through various high-throughput assays. The US100 D592 and D712 strains were characterized at sub-inhibitory concentrations through growth curves, RNA sequencing, bacterial cytological profiling, and exo-metabolomics high throughput experiments. The study reveals the vancomycin resistance behavior of USA100 strains in dual media conditions using wide-ranging experiments.

## Background and Summary

Methicillin resistant *Staphylococcus aureus* (MRSA) makes up the majority of hospital acquired *S. aureus* infections worldwide. Healthcare-associated MRSA (HA-MRSA) are subset of MRSA strains that often circulate in healthcare settings such as hospitals, dialysis centers etc. The MRSA infection is caused in healthcare settings like hospitals, dialysis centres, etc, thus often referred to as healthcare-associated MRSA (HA-MRSA) ^1,2,3^. The USA100 strain is the most common HA-MRSA with consistently high resistance to wide-range of antibiotics like macrolides, fluoroquinolones, and lincosamides ^4^. Moreover, they are considered to display vancomycin resistant and intermediate phenotypes ^5^. Vancomycin was first approved to treat MRSA in the late 1980s. However, by the 1990’s vancomycin-intermediate strains (VISA) had already begun to emerge. In 2002, the U.S. reported the first case of vancomycin-resistant *S. aureus* (VRSA) ^6^. To understand the genetic and phenotypic basis for the emergence of this resistance, we collected multi-omic data on HA-MRSA strains, D592 (daptomycin- and vancomycin-susceptible) and its descendent D712 (daptomycin-nonsusceptible and vancomycin-intermediate-resistant) previously collected from a patient with prolonged and persistent MRSA/VISA bacteremia for 21 days ^7^.

Cation-adjusted Mueller-Hinton broth (CAMHB) is standard medium for quantitative procedures for susceptibility testing in microbiology labs worldwide^8^. CAMHB is commonly used for antibiotic susceptibility as it is enriched in divalent ions, Ca^+2^ and Mg^+2^. The presence of divalent ions affects the stability of antibiotics or mode of action of antibiotics, which in turn greatly affect the minimal inhibitory concentration (MIC) values. Roswell Park Memorial Institute (RPMI) 1640 media is amongst the best media to mimic human physiology^9^.

The current study is focused on exploring the effect of vancomycin on HA-MRSA USA100 D712 and D592 strain in bacteriological (CA-MHB) and bacteriological (RPMI+10%LB) media. The MIC value of D592 decreased from 2**μ**g/ml R10LB to 1**μ**g/ml in CAMHB in presence of vancomycin Further, for D712 strain the MIC value decreased from 2**μ**g/ml in R10LB to 0.96**μ**g/ml in CAMHB. Here we interrogated the response of HA-MRSA strain to the sub-inhibitory concentration of vancomycin using growth curves, RNA sequencing, bacterial cytological profiling (BCP), and exo-metabolomics (HPLC and LC-MS). Together, our data provide an in-depth look into vancomycin response by simultaneously tracking gene expression (RNA-seq), cell morphology (BCP) and changes in the chemical composition of the media (HPLC and LC/MS).

## Methods

The methods are adapted from our previously published paper ^10,11,12^. The growth curve for the sub-inhibitory concentration is provided in **Figure 1**. The overall methodology is provided in **Figure 2**.

**Figure 1.**
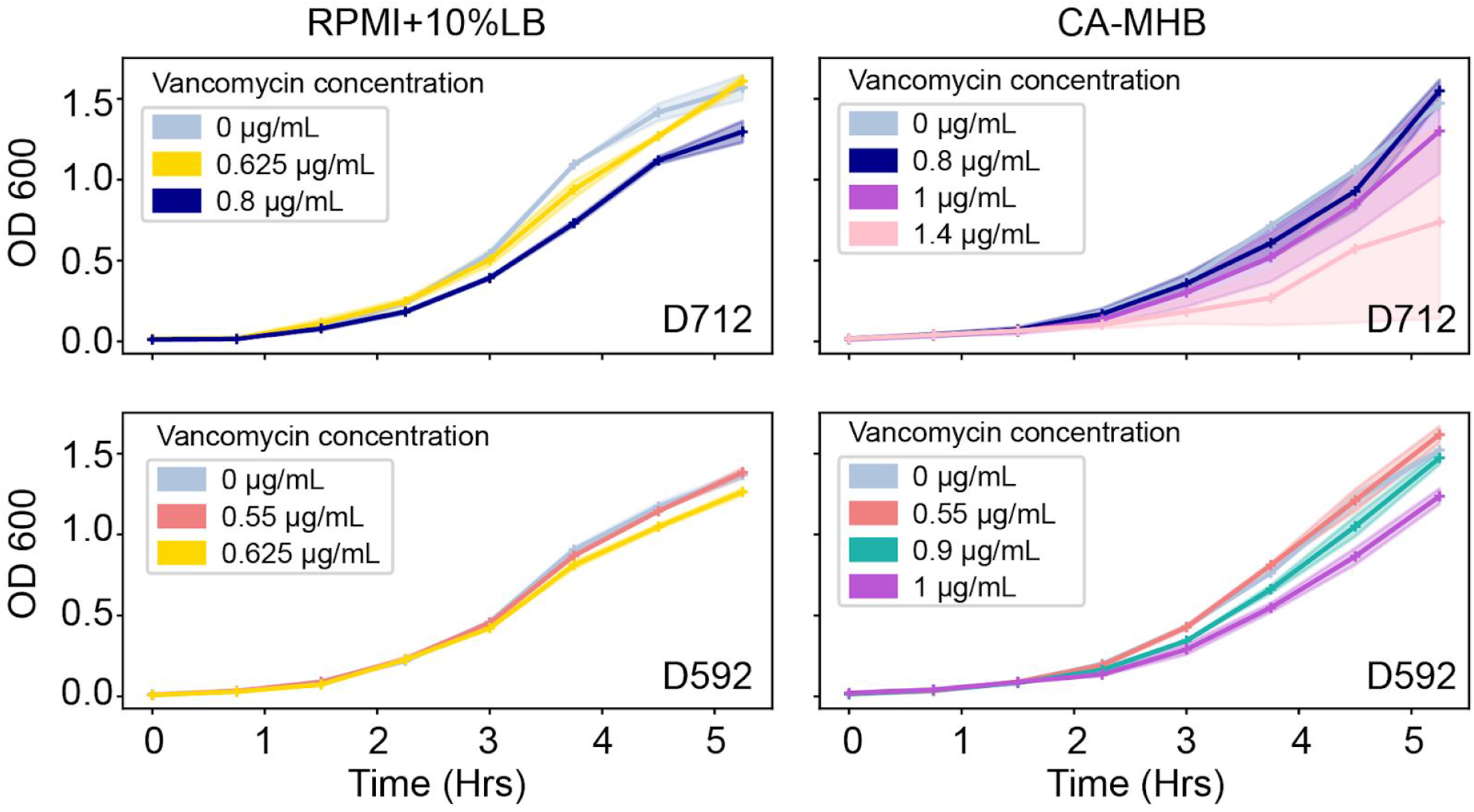
Growth curve for Staphylococcus aureus D592 and D712 strain in presence of vancomycin at various sub-inhibitory concentrations in CA-MHB and R10LB media.

**Figure 2.**
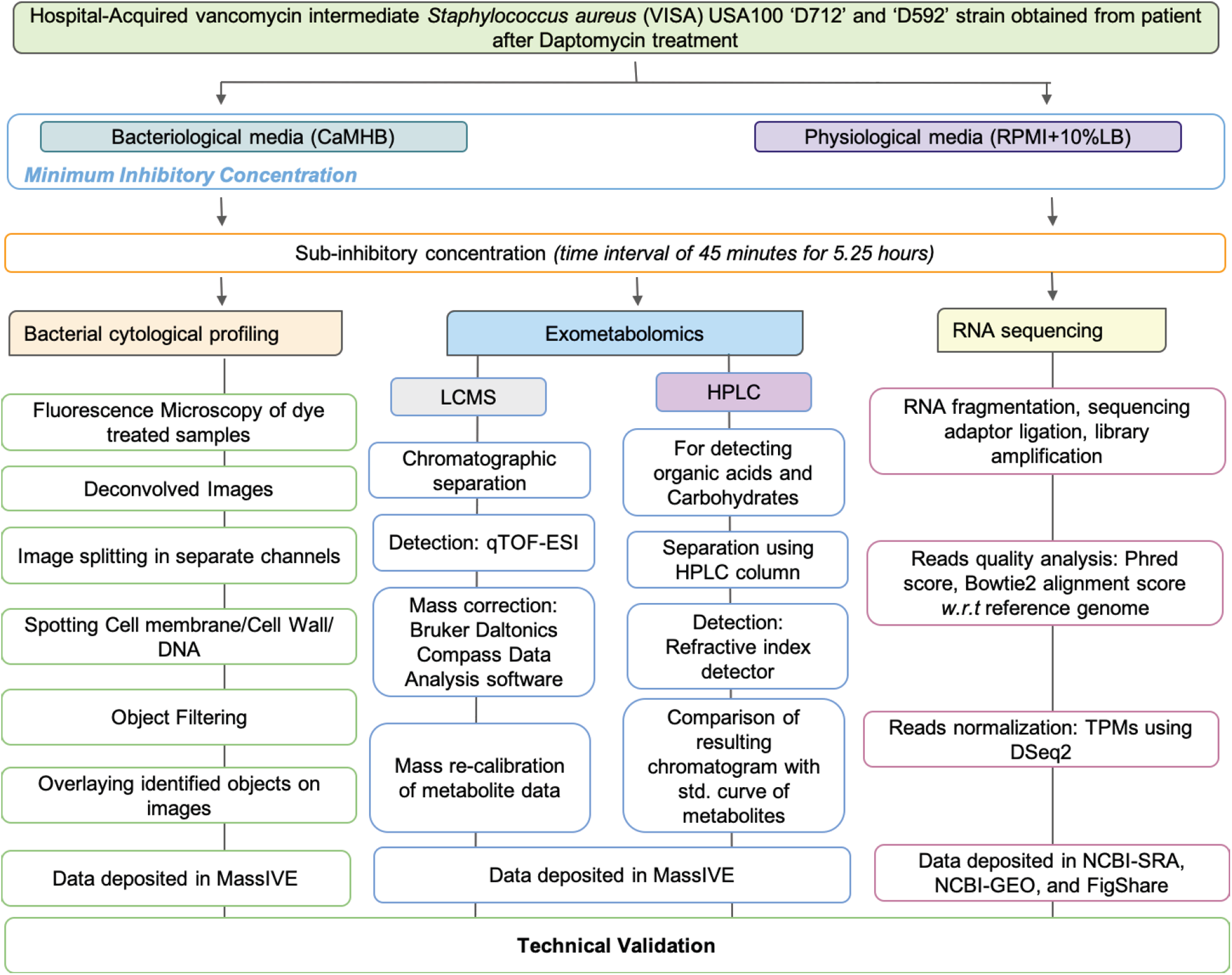
Diagram depicting the methodology of high throughput approaches used to profile the Staphylococcus aureus D592 and D712 in presence of vancomycin.

## Exclusion Criteria

The data of 1.4ug/mL sub-inhibitory concentration for CA-MHB on D712 strain has been excluded from all studies because the reproducibility between the samples was too low. The data and its result can be accessed in a public repository or available upon request.

## Data Records

The growth-rate data is available on Figshare^13^, while BCP, HPLC, and Mass spectrometry data have been deposited on MassIVE repository^14^. Complete RNAseq pipeline can be found at Figshare^15^, Fastq files of each run have been deposited on NCBI-Sequence Read Archive. More processed runs for RNAseq like TPM and counts can be found on NCBI-Gene Expression Omnibus. However, the overall summarized statistics of RNAseq is available on Figshare ^16^.

## Technical Validation

### Bacterial Cytological profiling

The technical validation of BCP was done through manual screening during the image segmentation process in CellProfiler. Accurate cell and object traces and measurements were verified manually for representative images. The cell outlines were matched with corresponding related structures e.g. DNA through “parent” tags. Finally, the output files for the cellular features were uploaded to MassIVE repository. A representation of the image analysis pipeline for BCP data is provided in **Figure 4**.

**Figure 3.**
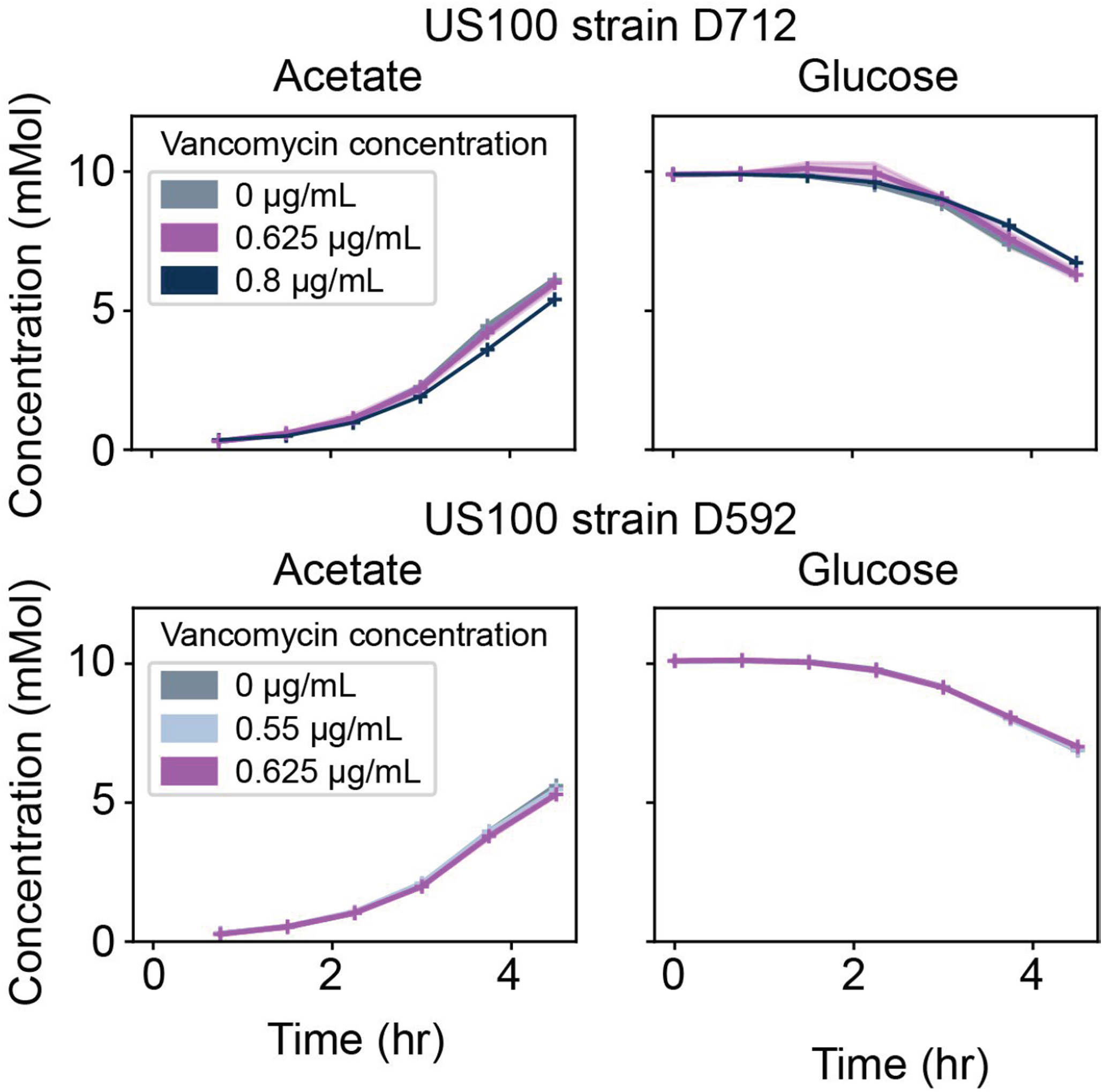
HPLC-derived quantitative time-course exo-metabolomics measurements for Staphylococcus aureus D592 and D712 cells exposed to various antibiotic concentrations in RPMI + 10%LB and CA-MHB. Here, we show the absolute calibrated concentrations of acetate and D-glucose in both media types, pyruvate in CA-MHB and lactate in RPMI + 10%LB.

**Figure 4.**
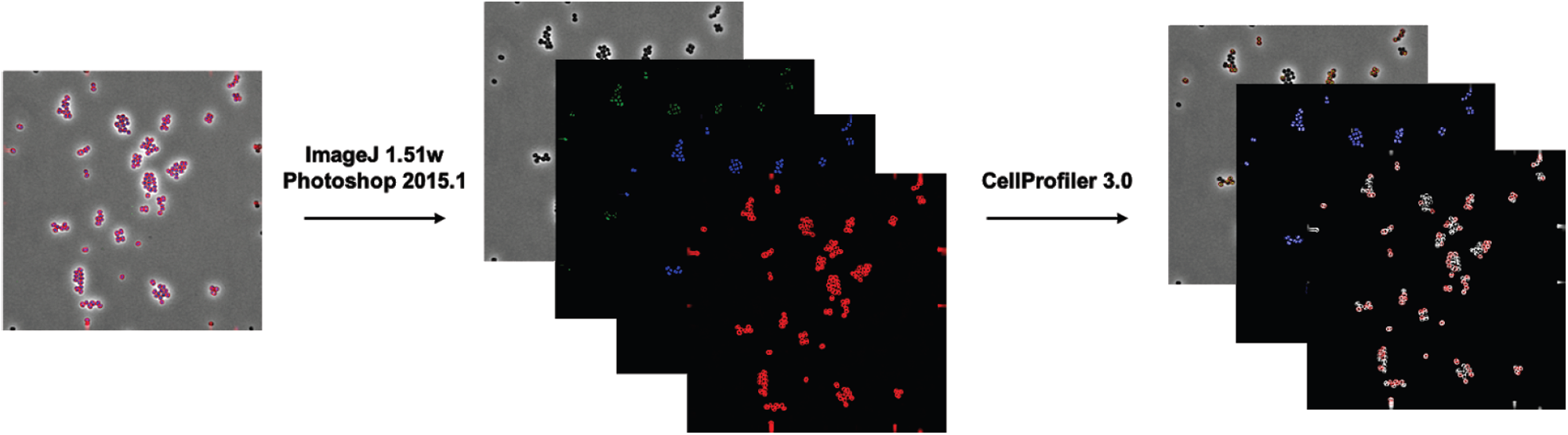
Depiction of image analysis pipeline for Bacterial cytological profiling of Staphylococcus aureus D592 and D712 in presence of vancomycin.

### Untargeted Liquid Chromatography Mass Spectrometry data acquisition

The reproducibility of global retention time and ion intensity were compared for each sample using the base peak chromatogram (BPCs) and multiple extracted ion chromatograms (EICs). The BPCs of each experimental triplicates were compared to get the reproducibility of retention time and peak intensity. While the EICs of the molecules were evaluated using retention time drift and peak area of <0.1 minutes and <15% respectively.

### RNA sequencing

The reference genome of D592 and D712 were sequenced using an Illumina Hiseq 4000. Prior to assembly, the quality control steps were performed to remove unincorporated primers, adaptors, and detectable PCR primers. The sequencing reads shows the average Phred score in D592 and D712 is >38.1 and >39.1 and respectively. The raw fastq files were uploaded at NCBI SRA web platform. The alignment of reads with the reference genome in D592 and D712 gives the alignment score of 98.55% and 99.52% correspondingly. The RNAseq results are shown in **Figure 5**.

**Figure 5.**
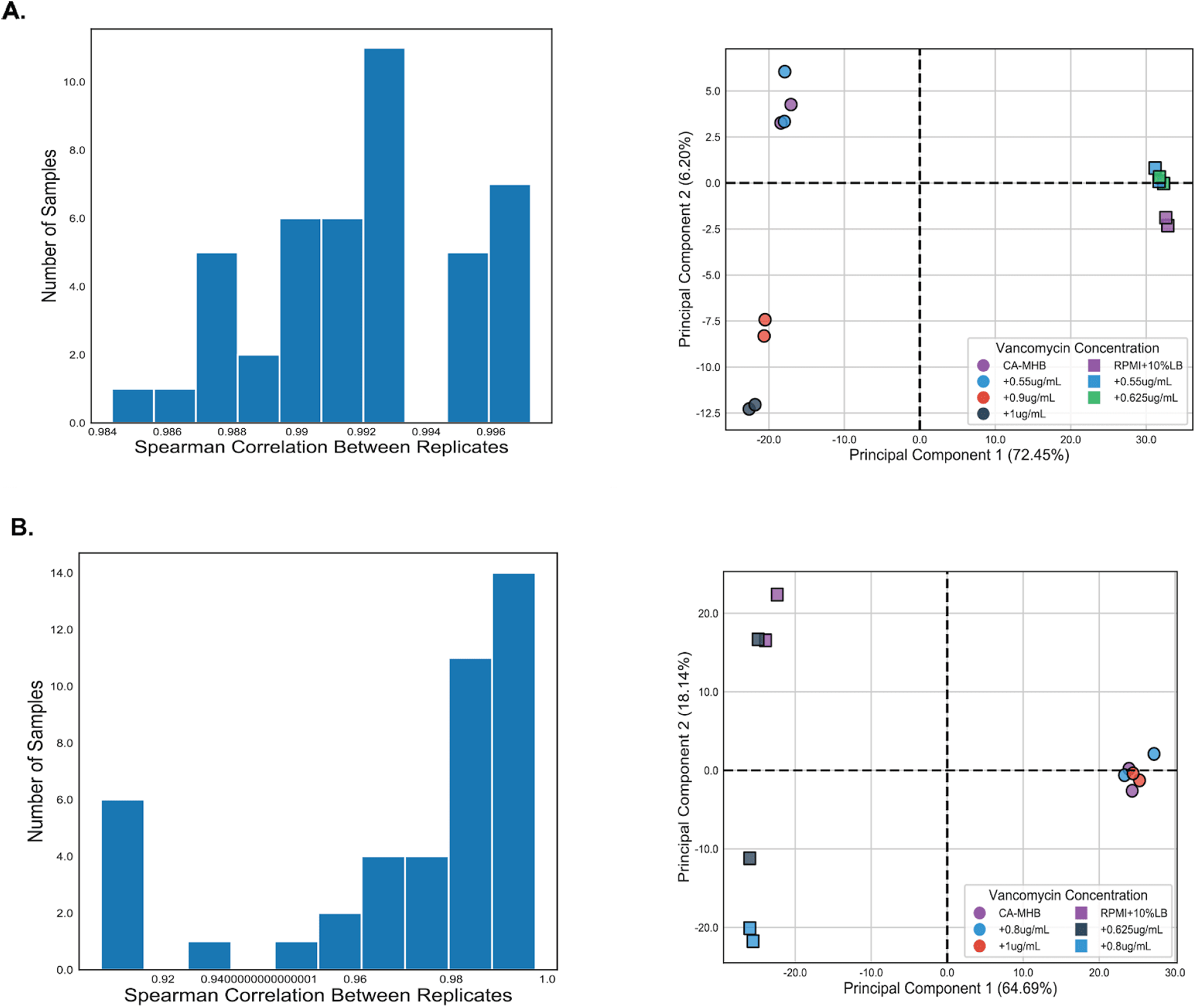
RNAseq results. A) Clustering of reads TPM as per Spearman’s correlation coefficient and PCA plot for D592 strains B) Clustering of reads TPM as per Spearman’s correlation coefficient and PCA plot for D712 strains.

## Code availability

The complete RNAseq pipeline used in analysis of RNAseq data is available on Figshare^15^.

## Acknowledgements

We thank Anand Sastry for helping build the RNA sequencing analysis pipeline. This research was supported by NIH NIAID grant (1-U01-AI124316).

## Author Contributions

A.R. compiled, analyzed results and wrote Data Descriptor, Methods and Technical Validation S.P. analyzed RNA sequencing data and wrote Methods H.T. performed growth experiments, analyzed BCP data, and wrote Methods M.M. analyzed HPLC data and wrote Data Descriptor, Methods R.S. prepared samples for RNA sequencing and wrote Methods C.A.O. prepared samples for HPLC and wrote Methods A.L. performed growth experiments, analyzed BCP data, and wrote Methods Y.S. analysed HPLC data. N.D. performed preliminary growth and MIC experiments. A.V. performed growth experiments. J.S. wrote Methods. S.M.D. performed growth experiments.

## Competing Interests

The authors declare no competing interests.

## Notes

### Competing Interest Statement

The authors have declared no competing interest.

